# Heat exposure and adverse pregnancy outcomes: evidence from humans and other viviparous species

**DOI:** 10.64898/2026.09.17.749897

**Authors:** Sofia Samoylova, Katherine A. Birchenall, Y. T. Eunice Lo, Sinead English

**Affiliations:** School of Biological Sciences, University of Bristol, Bristol, UK; Bristol Medical School, University of Bristol, Bristol, UK; School of Medicine and Public Health, University of Newcastle, Newcastle, Australia; Cabot Institute for the Environment, University of Bristol, Bristol, UK; Elizabeth Blackwell Institute for Health Research, University of Bristol, Bristol, UK

**Keywords:** heat exposure, birth outcomes, pregnancy, global warming, viviparous

## Abstract

**Introduction:** With global warming, heatwaves are increasing in severity, affecting species worldwide. Recent studies show associations between heat exposure in pregnancy and adverse birth outcomes such as preterm birth, low birthweight and stillbirth across multiple species. This suggests potentially conserved mechanisms such as maternal inflammation, dehydration and changes in placental physiology in response to heat exposure. Moreover, the gestational timing of antenatal heat likely impacts the type and severity of outcome, yet this is not fully understood.

**Methods:** We conducted a scoping review of studies on heat exposure and pregnancy outcomes in humans and other animals, to identify common mechanisms and temporal dynamics of effects. Temperature-, trimester- and species-specific patterns were reviewed across 197 studies and ten species (human, cow, sheep, pig, goat, rat, mouse, guinea pig, lizard and rabbit).

**Results:** Qualitative trends indicate that high temperatures are associated with adverse birth outcomes. In particular, third trimester exposure increases risk of preterm birth and low birthweight. Many studies emphasise placental involvement in adverse outcomes and point to potential aberrations in placental physiology or cortisol signaling pathways.

**Conclusions:** Overall, literature presents that heat exposure increases risk of adverse birth outcomes across studies, and taxa. Understanding the relationship between heat exposure and adverse birth outcomes, as well as the mechanisms and timing, is vital for informing public health policies and creating mitigating strategies for expecting mothers.

## 1. Introduction

There is mounting evidence linking heatwaves to increased public health risk via multiple biological processes (1–3). Pregnant women and their offspring are particularly susceptible to heatwaves due to the rapid metabolic changes and biological stress of pregnancy(4). Increased heat exposure has been associated with worse fetal and perinatal outcomes, including congenital anomalies, stillbirth, preterm birth, and low birthweight (Table 1; (5–9)). While there is an increasing body of evidence describing these associations, there remains limited understanding of the underlying mechanisms and possible temporal nature of these effects.

**Table 1.** Variables recorded from the studies.

| Topic | Subtopic | Variables collected |
| --- | --- | --- |
| **Study descriptors** | Study characteristics | Location, species, study type (cohort, time-series, case-control, observational, experimental) |
|  | Exposure timing | Fetal gestational age at exposure, method of gestational age |
|  | Environmental covariates | Humidity, air pollution |
|  | Temporal factors | Lag between heat exposure and outcome |
|  | Human-specific covariates | Maternal age, socioeconomic status, smoking status, urban vs rural residence, maternal ethnicity |
|  | Animal-specific covariates | Litter size |
| **Heat exposure** | Definitions | Temperature thresholds defining heat or heatwaves |
|  | Heat stress metrics | Heat stress indices (e.g., WBGT) |
|  | Temporal factors | Lag between heat exposure and outcome |
| **Outcomes** | Birth outcomes | Preterm birth, stillbirth, low birthweight, small for gestational age, PPROM |
|  | Critical window | Gestational window of susceptibility (if reported) |
| **Mechanisms** | Physiological | Placental weight, cotyledon number, placental volume, blood flow, nutrient supply, hypoxia |
|  | Genetic | Differential gene expression in cortisol pathway |
|  | Endocrine | Cortisol signalling, antidiuretic hormone, oxytocin |

Causal associations between heat and adverse birth outcomes are challenging to demonstrate, as potential effects associated with heat exposure may not always be immediately apparent(10): human gestation is relatively long, and the effects of a heatwave may be manifested weeks after exposure (11,12). Moreover, the timing of exposure during pregnancy is likely important, as per studies on other stress exposures showing critical windows(13,14). Heat exposure in late pregnancy has been associated with reduced gestational length, while early exposure is associated with an increase in frequency of babies born small for gestational age outcomes(15). In human cohorts, tracking prolonged exposure or exposure at various gestational stages is often challenging due to factors such as participant retention(16). Moreover, in prospective studies, participants are often enrolled after the first trimester, limiting insights on early exposures.

The effects of heat and heatwaves in pregnancy have been studied across several animal species. Although viviparity has evolved independently across multiple lineages, mechanisms such as hormonal regulation, plasma membrane transformation and egg retention have been conserved (17–20). Investigating heat effects in non-human viviparous models, where females of known conception date are tracked throughout pregnancy, and that have shorter gestational periods and higher fecundity, can lead to detailed mechanistic insights. These animal studies allow an alternative route to improve understanding of underlying physiological mechanisms, which may be conserved across species, due to the ability to do direct experiments and apply more invasive measures to investigate causality(21). It is, however, vital to consider differences among species such as placental characteristics, gestation length and thermoregulation mechanisms, when comparing any mechanisms among species, including links to human health(18).

While species differ in placental physiology, this structure remains an essential organ during gestation that may mediate heat induced pregnancy outcomes, as it functions to support the growing fetus with nutrition, endocrine processing, waste elimination and other functions(22). Placental weight according to gestation is a key physiological factor, often used as a development and function proxy for the placenta(23). Decreased placental weight for gestation is associated with adverse outcomes such as placental abruption, preeclampsia and fetal growth restriction(24–26). Several studies have noted a heat-mediated placental role in adverse birth outcomes such as low birthweight and stillbirth, through a range of mechanisms such as placental blood flow or the initiation of inflammation pathways(27,28). The exact pathways and genes involved, however, are understudied and require further elucidation.

To our knowledge, previous scoping and systematic reviews investigating the association between extreme heat and birth outcomes(5,9,29) have focused on humans only. Here,–we specifically investigate the potential of conserved patterns and mechanisms in birth outcomes across animals, and with a particular focus on the temporal dynamics of heat exposure. Additionally, we expand on existing methodologies by outlining the variation in organism-specific and general heatwave definitions for studying heat-related birth outcomes and discussing whether a consensus measure would be helpful. By taking this comparative taxonomic perspective and focusing on mechanism and temporal dynamics, this review aims to identify existing gaps in the literature, providing a future direction for research.

## 2. Methods

### 2.1 Protocol and registration

A review protocol has been created and uploaded on OSF on December 22, 2023. The protocol can be accessed on: https://osf.io/wa4tc/wiki/home/.

### 2.2 Data sources

An initial search in SCOPUS, PubMed and Google Scholar was undertaken in October 2023 to determine the databases with the highest result number and identify additional exclusion key words. The main search included terms relevant to high temperatures and climate change, and terms relevant to birth outcomes. Google Scholar was excluded due to results from SCOPUS and PubMed being sufficiently comprehensive. Search results returned 5764 publications in SCOPUS (in October 2023) and 4164 publications in PubMed (in December 2023). Initial screening of all results included elimination by title, followed by elimination by abstract and contents.

Due to the increasing research in the field, a secondary search was performed in February 2026, yielding 743 publications in Scopus and 582 publications in PubMed. Forty-one additional publications were added to the data collection after paper elimination.

### 2.3 Search terms

We selected the following inclusion terms due to their relevance to the research topic: TITLE-ABS-KEY ( (“heat” OR “heatwave” OR “heat event” OR “elevated ambient temperature” OR “climate change” OR “urbanization” OR “heat islands” OR “high temperature” OR “warm temperature“) AND (“pregnancy” OR “birth” OR “preterm” OR “preterm birth rate” OR “gestation” OR “maternal” OR “low birthweight” OR “stillbirth“).

In addition, exclusion criteria were selected to eliminate non-viviparous organisms, lactation-related outcomes, common non-temperature environmental factors, paternal factors, pre-existing maternal conditions, genetic disorders and non-relevant terms that appeared upon preliminary searches (Supplementary Table 2).

### 2.4 Eligibility criteria

Publications reviewed include mothers of various viviparous species (animals that develop the embryo inside the mother’s body) that have experienced elevated temperatures or heat stress events and their birth outcomes. Publications were excluded if they provided a vague term such as “heat” and did not provide specific temperature values (e.g. median temperature) or a quantitative definition of what was considered heat stress. Publications were also excluded if there was no full text access. There was no inclusion criteria based on publication date as the field was relatively new, the review aimed to collect all literature available and consider all methodology and mechanisms suggested. The heat exposure included both exposure up to three months pre-conception and during the gestational period. Birth outcomes included stillbirth, low birthweight, and preterm birthweights (Table 1). We did not apply a cut-off age limit for females, as we were interested in effects across the reproductive lifespan. Only English-language publications were included.

### 2.5 Data Synthesis

Data from each study was tabulated (Table 1), including a range of maternal physiological, environmental and socio-demographic variables known or likely to be associated with responses to heat exposure. Respective risk ratios and 95% confidence intervals as reported by authors were also collected for outcomes (Definitions present in Table 2).

**Table 2.** Included adverse birth outcomes and their definitions.

| **Term** | **Definition and potential underlying mechanisms** | **Evidence of heat-exposure association** |
| --- | --- | --- |
| **Preterm births** | Births occurring prior to thirty-seven completed weeks of gestation (i.e. <37+0), with further subclassifications into spontaneous, preterm prelabour rupture of membranes (PPROM, see below) and medically induced preterm birth(9,30). | [S4] [S5] |
| **Preterm Prelabour Rupture of Membranes (PPROM)** | Rupture of fetal membranes prior to 37 weeks' gestation, and before onset of labour; cause often unclear, but can be due to intrauterine infections or overdistention(32,33). | [S6] [S7] |
| **Stillbirth (also known as intrauterine fetal demise) and miscarriages** | The definition of stillbirth and miscarriage varies between countries and different organisations (34), but stillbirth generally is defined as a pregnancy loss after 20 to 28 weeks' gestation, although some classifications use birthweight (i.e. deliveries >400g or >500g resulting in death)(35). One of the included studies defined stillbirth as death after 7 months [S48]. Miscarriage definition also differs geographically and will align with stillbirth definitions: for example, in countries where the definition of stillbirth is defined as pregnancy loss after 20 weeks, miscarriages will refer to loss prior to 20 weeks. While stillbirth and miscarriages differ in definition, they are often grouped in studies due to shared risk factors and resulting in a loss of pregnancy as well as changes in gestation of viability with the improvement of neonatal care(36,37). | [S9] [S1] (S10)(39) |
| **Low Birthweight** | Birthweight less than 2500g at term(40). | [S11] [S12] |
| **Small for gestational age (SGA)** | Birthweight less than the 10<sup>th</sup> centile for gestation. While not fully overlapping with the low-birth-weight definition, SGA is closely related as a birth outcome and often results in similar health prognoses(41). Some studies therefore group these outcomes together when analysing environmental effects, such as temperature variations, on birthweight outcomes (9). | [S13] |

## 3. Results

Searches from SCOPUS and PubMed produced 11253 publications (10796 post duplicate removal), with subsequent elimination according to eligibility criteria on screening titles, abstracts, contents resulting in 197 publications (Figure 1A). These studies included the following outcomes: 88 human studies and 11 animal studies analysing preterm birth, 6 human studies on PPROM, 31 human studies and 18 animal studies analysing stillbirth or miscarriage, and 50 human and 32 animal studies analysing birthweight (Supplementary Table 3). Some papers are included in more than one category because they analysed multiple birth outcomes.

**Figure 1.**
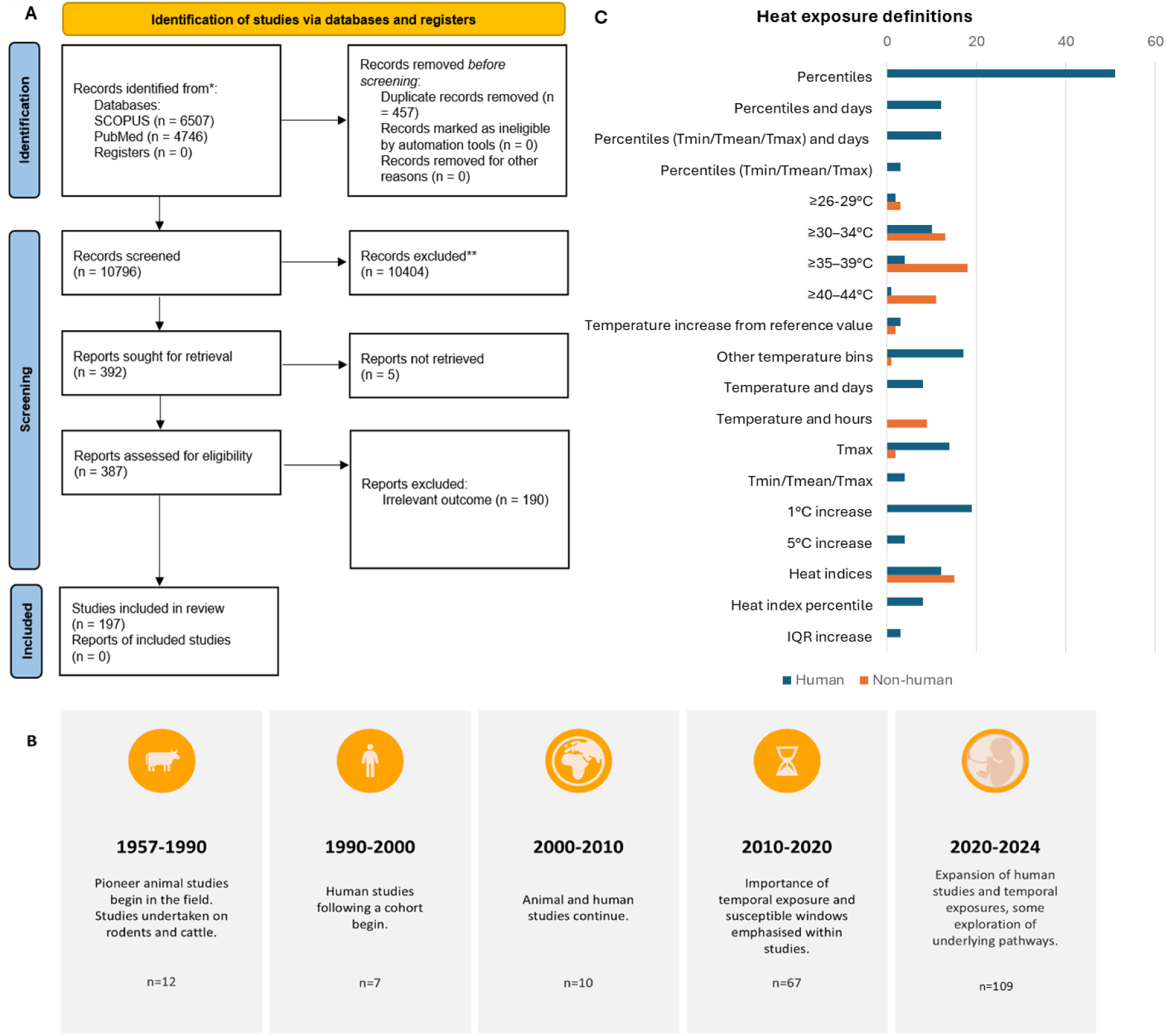
Data synthesis, paper timeline and heat stress definitions. **1a)** PRISMA flowchart showcasing the selection and elimination processes for the publications. 11253 publications were originally identified from SCOPUS and PubMed, with subsequent elimination of 457 duplicates. Upon title and abstract review 10404 papers were removed, with a further 189 papers eliminated once contents were analysed, resulting in 197 publications in the final list. **1b)** Approximate timeline and patterns observed of included studies., showing the number (n) of studies per time window. **1C)** Frequency of heatwave definitions in human (blue) or non-human animal (orange) studies. Percentiles = percentage points showing mean temperature distribution above a certain value stress. Percentiles _min/mean/max_ = percentage points showing minimum, mean and maximum temperature distributions above a certain value.) Percentiles and day = percentage points showing mean temperature distribution above a certain value stress over the span of several days (usually 2-5 depending on study). Temperature and days = a select temperature point selected as a heatwave value over the span of several days (i.e. >32C across 2 days) T_mean_/T_max_/T_min_ = mean, maximum and minimum temperature as some studies utilise all three values to determine heat stress. 1C (or 5C) increase = 1 degree (or 5 degree) increase increment (s) from median/reference value. Indices = exact value of indices such as heat index, wet bulb globe temperature, temperature heat index, apparent temperature, Universal Thermal Climate Index. Indices and percentiles = percentile value of indices. Definitions utilised by less than 3 studies were omitted but are present in the supplementary information.

Macfarlane et al., 1957 and Edwards, 1969 (Table S1) were among the first to consider heat stress effects on birth outcomes, in mice and cattle. From 1990, the field shifted towards human studies, with the number of publications notably increasing since 2010 (Figure 1B). Of the studies included in this review, 168 were published after 2010. These more recent studies had an increased emphasis on temporal heat exposure, susceptible windows during pregnancy, and lagged effects on birth outcomes.

There is no universal temperature threshold for heat or a heatwave because what is considered elevated temperature depends on local climate, temperature variability, species of interest, and their capacity to cope with heat (43) . The included studies used a variety of temperature or heat stress thresholds (Figure 1C). The most common approach (65 studies) used temperature bins (i.e., 30-35°C, 35-40°C), followed by temperature percentiles, usually the 90 or 95^th^ percentiles of local temperature distribution (49 studies). Multiple studies relied on one degree increases relative to the mean. Thirty-nine studies used heat stress metrics such as wet-bulb globe temperature (WBGT) and Heat Index (HI). WBGT and HI incorporate both temperature and humidity, based on different thermal comfort algorithms(44). Fifty-two studies utilised multiple heat exposure definitions.

Geographically, 41 studies were in USA, 33 in China, 20 in Australia, 8 in Israel, Uganda and Spain, 7 in Malawi and 6 in Ghana, India, Kenya, Nigeria, South Africa, South Korea and Zimbabwe (Figure 2A). Other locations had fewer than 5 studies, with significant gaps in Eastern Europe, Northern Africa, Southeast Asia and South America. Studies covered 10 species (Table S3), with 147 human, 14 rodent, 13 bovine, 11 ovine, 4 porcine, 4 reptile, 3 caprine and 1 rabbit studies (Figure 2B). Most animal studies (37 out of 50) had an experimental design. For confounding and additional factors, 82 studies considered socio-economic status or a relevant proxy, 81 effects of maternal age, 77 humidity effects irrespective of temperature, 68 air pollution, 45 maternal ethnicity, 39 cigarette smoke (including second-hand exposure) and 24 urban vs rural settlement (Citations in Supplementary Table 3).

**Figure 2.**
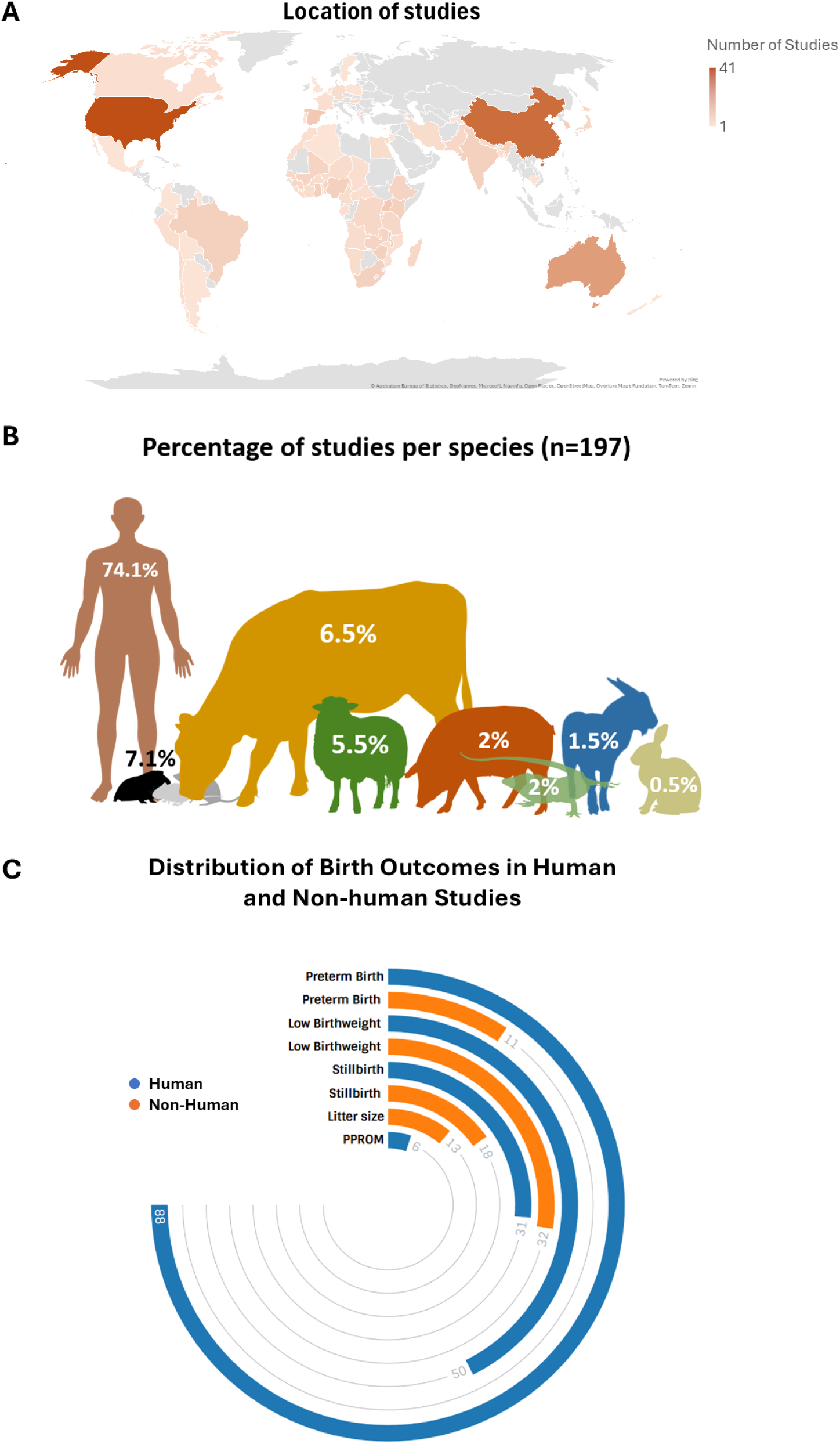
Birth outcome distribution, geographical locations, species distribution of the 197 studies included. 2A) Geographical distribution of studies collected for the review. 2B) Percentage representation of the different animal species included in order of appearance (human, rat, mouse, guinea pig, cow, sheep, pig, lizard, goat and rabbit) and their distribution. 2C) Radial bar chart showing the distribution of birth outcomes between human (blue) and non-human (orange) by studies. Count of studies indicating each outcome is indicated by number at the end of the bar.

### 3.1 Preterm births

Of 88 human studies including preterm birth as an outcome, 80 found an increased risk of pre-term birth with heat exposure (Figure 2C; risk ratios, RR, from 1.004 (95% Confidence Interval (1.001–1.007) to 3.35 (1.39 - 8.06)). Six studies found a higher temperature associated with reduced preterm birth risk: one presented reduced risk when analysing subgroups of maternal ethnicity, one when utilising a specific heat model (Mean daily temperature > 99th percentile for ≥ 2 consecutive days) and two when specific exposure timings such as preconception or whole pregnancy exposure was analysed. Five studies showed no statistically significant associations.

Specific windows of heat exposure during gestation were investigated in 73 studies, with increased association or incidence of preterm birth following third trimester exposure (Figure 3A). Lag effects were considered in 36 studies with the most common outcomes showing short windows of 0-7 days or a lag of one month for pre-term birth.

**Figure 3.**
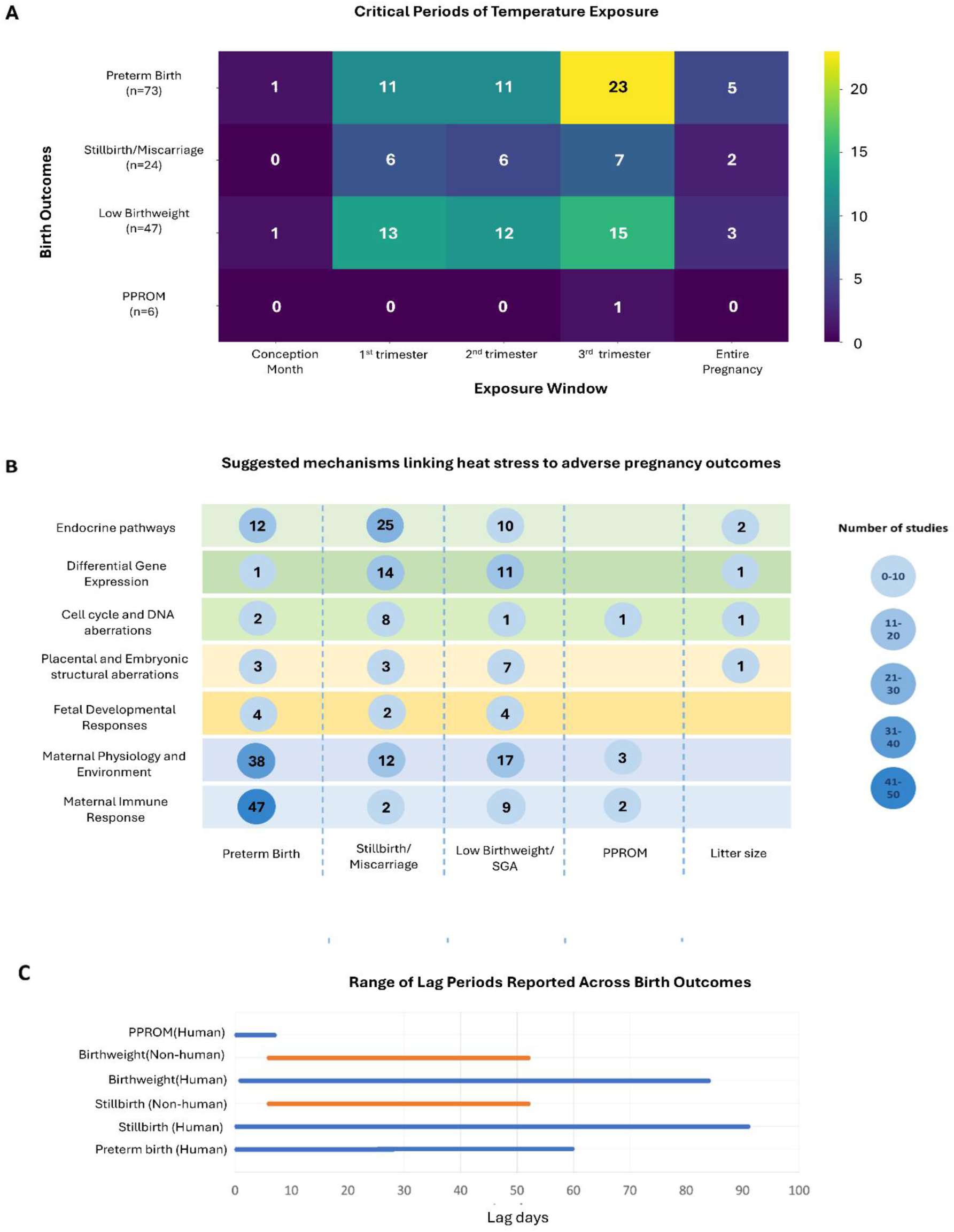
Temporal effects, physiological mechanisms and lag effects of heat exposure in pregnancy with outcomes. A) Temporal effects in human studies which consider exposure across pregnancy (n=124 total). Number of studies written in box and shading denotes the number of studies which show an effect within a specific trimester. The critical period of exposure in this review is defined as the period that a study has reported a significantly elevated risk ratio or coefficient for the adverse pregnancy outcomes. Some papers are included in more than one category: the same paper could analyse multiple birth outcomes and therefore listed as multiple counts. **B)** Suggested mechanisms include altered endocrine signalling, disruption of cellular processes, changes in maternal physiology, disruption of the immune landscape and disruption in placental development. **C)** Range of significant lag periods in days reported in studies across birth outcomes between human and non-human studies.

In animals, heat exposure is associated with reduced gestational length (10 out of 11 studies). However, one study did not find statistically significant results, and one rodent study showed a 4% increase in gestational length. No lag effects were analysed.

### 3.2 PPROM

Six studies considered PPROM, including two ecological observational studies, three cohort and one case-crossover study, and all showed a positive relationship between temperature and PPROM risk (RR range: 1.081 (1.005–1.160) - 2.161 (1.240–3.764)). Lag times varied from immediate to a one-week delay.

### 3.3 Stillbirth and miscarriage

Thirty-one human studies investigated heat effects on stillbirth and miscarriage (Supplementary Table 3); with 27 showing a positive correlation between temperature and stillbirth risk (RR from 1.012 (1.010-1.013) to 3.81 (3.69-3.94)). Overall, the included studies did not identify a distinct pattern regarding a critical window of exposure. One study found a protective effect for stillbirth with first trimester exposure (RR 0.73 (0.58 – 0.93)). Lag effects varied from immediate to three months.

Eighteen non-human studies measured prenatal mortality, with 14 finding a decrease in embryo viability or an increase in stillbirth rates associated with heat exposure (Supplementary Table 3). Eight studies were performed on rodents and included a range of developmental stages, with six finding embryo loss at earlier developmental stages such as the morula and blastocyst. The increased risk ranged from 1.21 to 1.83, depending on gestational stage, compared to background rates ranging from 1.03 to 1.27.

### 3.4 Low birthweight, small for gestational age, and decrease in birthweight

Heat exposure on low birthweight was measured in 50 human studies, with 45 finding a negative correlation between elevated temperature and birthweight. Studies also largely utilised percentile thresholds, most commonly with temperatures over the 90^th^ or 95^th^ percentile. The outcomes within studies varied with the weight decrease ranging from −54.2 g (−102g to -6g) to -287.4 g (−474.1g to - 100.8g) for 1°C increments. Risk ratios for low-birthweight ranged from 1.054 (1.016–1.094) for the 90^th^ centile compared to the mean, to 2.89 (1.17 - 7.14) for the 95^th^ centile compared to the mean. SGA risk ratios presented by the studies ranged from 1.041 (1.029-1.054) when the 90^th^ percentile was compared to 40th to 50th percentiles, to 4.7 (1.6-14.1) for 95^th^ centile comparison to the 5–95th percentile. Three studies had no statistically significant outcome. While the reported critical windows varied across studies, 15 studies found exposure in the third trimester to be particularly significant. Four studies presented lags with values ranging from a week to 14 weeks.

Thirty-two non-human studies measured offspring weight, with 23 showing a decrease in weight with heat exposure.

### 3.5 Litter size

Thirteen non-human studies looked at the effects of heat exposure and temperature variations on litter size, with 6 showing a decrease in litter size with high temperature exposure, 6 no effect, and 1 an increase in litter size. While the studies included rodent, rabbit, reptile, and pigs, most (54%) were on rodents.

### 3.6 Suggested mechanisms

Fourteen studies looked at potential underlying mechanisms, with only two on humans, which emphasised the role of placental physiology showing a decrease in placental weight, volume and blood flow with heat exposure. Ten studies looked at physiological mechanisms, with nine presenting a decrease in placental weight with heat exposure. Other mechanisms considered in single studies include decrease in blood flow, glucose and oxygen uptake, differential gene expression, and upregulation of antidiuretic hormone and oxytocin. Some suggested but unexplored mechanisms within these studies have been listed in Figure 3B.

Placental physiological factors have been linked to heat in multiple species(45,46), however, causative relationships – particularly in humans – have not been clarified thus calling for further studies.

## 4 Discussion

### 4.1 Overview

We reviewed the literature to understand effects of heat exposure on adverse birth outcomes in humans and other animals, focusing on temporal effects and underlying mechanisms. Most studies found an association between heat exposure and an increased risk of adverse birth outcomes, but a small number of studies (9 out of 197) noted a potential protective relationship. There was a potential sensitive period of heat exposure in third trimester linked to preterm birth and low birthweight; however, no overall sensitive periods appeared to be present for the other outcomes. Most non-human animal studies took an experimental approach, finding lower birthweights in heat stressed groups compared to controls. Mechanistic studies – the majority being in animals – largely focused on heat exposure effects on physiological processes involving the placenta, including placental weight, blood flow and nutrient transfer, which in turn affect could fetal growth and viability.

### 4.2 Temporal effects: Critical windows and lag effects

We analysed patterns in 124 of the human studies which compared outcomes after exposure throughout the entire gestational period. Of these, 45 studies presented an elevated risk in adverse birth outcomes linked to third-trimester exposure: 23 studies showed an increase in preterm birth risk, while 15 studies showed a decrease in birthweight (Figure 3). This pattern is not observed in previous systematic reviews on heat exposure and pre-term birth. For instance, Chersich et al., (2020) and Lakhani et al., (2024) either found an increase in risk across gestational periods or that the first and second trimesters were critical periods of exposure(5,47). This could be due to differences in study numbers: our review consists of 197 studies while Chersich and Lakhani had 70 and 16, respectively; and it is also worth noting ours focuses on qualitative patterns rather than being a formal meta-analysis. Furthermore, specific identification and establishment of casual associations of critical windows can be constrained due to time lags, as the effect could be perceived to be associated with other variables rather than heat exposure.

Out of 50 animal studies only 11 reported any significant critical windows . Results varied depending on species, with reptile studies reporting exposure over the entire pregnancy to be significant for preterm birth whereas mice tended to show an increase in adverse birth outcomes such as low birthweight and miscarriages with exposures earlier in the pregnancy. With these inconsistencies, the need for further investigation of critical windows such as further longitudinal studies with larger sample sizes is highlighted, which is further supported by the fact that only just over a third of the included studies consider temporal windows. Increasing evidence from human studies suggests that timing of stress exposure can be vital for pregnancy outcomes, with historical cohorts such as the Dutch Famine birth cohorts demonstrating that birth outcomes depended on exposure timing, with earlier exposure being linked to metabolic syndrome while mid to later exposure was linked to increased frequency in depressive disorders (48–50).

Sixty-five studies considered lag effects, but no common lag time was found. The most common approach to analysing lag effects among these studies included analysing specific cumulative lag values (e.g., lag 0–6 days), closely followed by the approach of using linear distributed lag models. Effects varied from being evident immediately (lag 0) to presenting after 14 weeks regardless of the birth outcome (Figure 3C). The differences in lag values between birth outcomes are likely due to differences in underlying mechanisms(11), while variation in results for the same birth outcomes could occur due to study location and environmental factors, or population differences such as socioeconomic status and thus difference in access to heat mitigating actions(5). A systematic approach with model adjustment for various confounding factors and population differences could aid in standardising results, thus providing a more accurate analysis of potential lag times.

### 4.3 Proposed mechanisms across animal species

Across species, reviews such as Laporta et al. (2024), Wyrwoll (2023), and Hansen (2009) highlighted similarities in heat-related adverse birth outcomes, although there was some bias in livestock studies towards analysing birthweight(51–53). These adverse birth outcomes were thought to likely stem from placental function changes.

A small subset of studies, predominantly in animal models, report decreased placental weight, blood flow, oxygen and glucose uptake, and decreased placental cotyledon size in relation to heat. The placenta is a vital organ responsible for fetal blood, oxygen, hormone and nutrient supplementation as well as waste removal. Any placental aberrations therefore directly influence the fetus and can lead to disruption in development (54).

There are however some limitations in interspecies comparison which lie in differences in types of pregnancy, in terms of gestation duration, extent of placentation, and litter size (55,56). Moreover, thermal physiology varies, for example both endothermic and ectothermic species were included which may complicate comparison(57). Interestingly, the ectotherm studies on lizards (n=4; [S81][S84][S85][S141]) seemed to mimic patterns observed in endothermic viviparous organisms, with a decrease in gestational length with heat stress. Another challenge lies in placental physiology differences, both between species and when extrapolating results to humans (19). While some mechanisms and endocrine pathways are conserved among species (58–60), future comparative research should account for differences in placental physiology. Additionally, animal studies often follow an experimental design as opposed to observational, making it vital to increase observational animal studies in field conditions, to be able to observe more natural effects of heat on viviparous birth outcomes.

While mechanisms related to placental physiology and function have received some attention(29,54), less is known about non-placental mechanisms. Additional heat-stress associated mechanisms (Table 2) suggested to cause adverse pregnancy include decreased blood flow to the fetus due to blood flow diversion to maternal skin, upregulation of prostaglandin and oxytocin, inflammation, increased blood coagulation uterine contractions(61), and malformations that result in stillbirth or miscarriage. These mechanisms could contribute to the increase in preterm birth risk in the third trimester. Genetic factors have also had comparatively less attention, with more work required in this area. This, however, could be reflective of the limitations of our search scope concentrating on heat exposure and physiological outcomes rather than molecular studies as recent studies have shown an association between early environment and offspring DNA methylation(62).

Despite an increase in studies linking heat and adverse pregnancy outcomes, knowledge gaps remain. This is highlighted by the lack of studies in our review that have included placental analysis: 14 out of the 197 studies considered potential underlying mechanisms, with only 2 human studies. Factors such as heat-mediated changes in gene expression – such as cortisol and oxytocin – are mentioned but remain largely unexplored(63). Studies such as Lakhani et al. (2024) have also suggested considering biomarkers that may indicate a mother is more susceptible to heat-mediated effects on birth outcomes. This ranges from the presence of inflammatory markers such as interleukin – 6 to looking at the levels of vasodilators ,which would help the mother regulate her blood flow in response to heat stress, such as nitric oxide(64,65). This could It is important to consider these factors as they give insight into the pathways linking heat stress and pregnancy outcomes and could help inform mitigation strategy. Their omission in previous research could be due to several factors including the requirement for additional ethical agreements when obtaining biological samples, storage and sequencing costs, and historically limited consideration of female biology in research(66,67). Future cohort studies should increase the emphasis on placental monitoring and aim to determine underlying mechanisms. Birth cohorts such as The Avon Longitudinal Study of Parents and Children and The New Hampshire Cohort study now collect and analyse placental tissue, allowing the observation of physiological and transcriptomic data(68,69). Moreover, further comparative transcriptomic and DNA methylation analysis – both on placenta and cord tissue – could help identify the conserved networks involved (70,71).

### 4.4 Heatwave definitions

Heatwave definitions varied between studies and organisms, with human studies largely utilising percentile-based definitions, while non-human studies considered heat indices in observational studies and set temperature points in the case of experimental studies.

Differences in heat definitions and measurements may have arisen due to study design. While observational and experimental studies may analyse the same temperature point, experimental studies – especially with climate-controlled chambers – provide the advantage of more stable conditions(54). Standardisation of study designs, or model adjustments should therefore be considered in future studies. Considering the impact of global warming on ambient temperatures, percentile reference periods should be considered – the definition of a temperature percentile for the same location may vary depending on the time period(72). Additionally, definitions such as 1C increments over baseline can vary depending on geographic context as the same increment will result in different ambient temperatures and thus physiological responses. While studies have looked at temperature variation within a week or month, diurnal variations – specifically elevated night-time temperatures – have been widely overlooked. Diurnal temperature variations have been shown to influence sleep and thermoregulation which may in turn affect cardiac and respiratory functions, highlighting their potential relevance for future analysis (73–76). As the global climate warms, heatwave definitions may continue to change. It is therefore vital to refine these definitions to ensure they remain relevant and effective in determining heat-related risks.

## 5 Conclusions

This review brings a comparative perspective to understand the association between heat exposure and adverse birth outcomes across humans and other animals, considering the mechanisms and temporal dynamics. Our review of the literature suggests that the mechanisms may largely rely on placental function and hormonal signalling, calling for further studies to be undertaken with the consideration of these factors. It additionally draws attention to a potential critical exposure window for preterm birth in the third trimester, which has not been fully observed in previous studies.

Despite a growing number of studies in this area, several gaps remain, calling for further genetic analysis and minimising the heterogeneity of study designs. Developing a more standardised approach to studies, both in terms of heat definitions and in study design and critical windows considered, will allow to produce more comparable results. As global temperatures continue to rise, a deeper understanding of heat exposure effects on pregnancy outcomes across different populations and species will be crucial in developing comprehensive strategies to address climate-related reproductive health challenges.

## Abbreviations

WBGT: Wet bulb globe temperature
PPROM: premature rupture of membranes
HI: Heat Index
SGA: Small for gestational age
RR: risk ratios

## Statements and declarations

### Author Contributions: CRediT

**Sofia Samoylova:** Conceptualization, Methodology, Investigation, Data Curation, Writing - Original Draft , Visualization, Project administration, Formal analysis ; **Katherine A. Birchenall:** Conceptualization, Methodology, Funding acquisition , Writing - Review & Editing , Visualization, Supervision, Project administration; **Y.T. Eunice Lo:** Conceptualization, Methodology, Funding acquisition , Writing - Review & Editing , Visualization, Supervision, Project administration; **Sinead English:** Conceptualization, Methodology, Funding acquisition , Writing - Review & Editing , Visualization, Supervision, Project administration

### Data access statement

The dataset generated can be accessed at: https://osf.io/wa4tc/files/x2twv. Information about the publications included such as study design and locations can be found in the supplementary files.

### Ethics approval and consent to participate

Not applicable

### Consent for publication

Not applicable

### Authors contribution

SS led data extraction, analysis, and manuscript drafting. YTEL, KB, and SE contributed to study design, critical revision of the manuscript, and interpretation of the findings. All authors approve the final version of the manuscript. SS is the guarantor.

